# Spatial transcriptomics reveals distinct and conserved tumor core and edge architectures that predict survival and targeted therapy response

**DOI:** 10.1101/2022.09.04.505581

**Authors:** Rohit Arora, Christian Cao, Mehul Kumar, Sarthak Sinha, Ayan Chanda, Reid McNeil, Divya Samuel, Rahul K. Arora, T. Wayne Matthew, Shamir Chandarana, Robert Hart, Joseph C. Dort, Jeff Biernaskie, Paola Neri, Martin D. Hyrcza, Pinaki Bose

**Author notes:** Pinaki Bose PhD. Room 354, Heritage Medical Research Building, 3330 Hospital Drive NW, Calgary, Alberta, Canada T2N 4N1. These authors contributed equally (Rohit Arora, Christian Cao).

## Abstract

We performed the first integrative single-cell and spatial transcriptomic analysis on HPV-negative oral squamous cell carcinoma (OSCC) to comprehensively characterize tumor core (TC) and leading edge (LE) transcriptional architectures. We show that the TC and LE are characterized by unique transcriptional profiles, cellular compositions, and ligand-receptor interactions. We demonstrate that LE regions are conserved across multiple cancers while TC states are more tissue specific. Additionally, we found our LE gene signature is associated with worse clinical outcomes while the TC gene signature is associated with improved prognosis across multiple cancer types. Finally, using an *in silico* modeling approach, we describe spatially-regulated patterns of cell development in OSCC that are predictably associated with drug response. Our work provides pan-cancer insights into TC and LE biologies, a platform for data exploration (http://www.pboselab.ca/spatial_OSCC/) and is foundational for developing novel targeted therapies.

## Background

Oral squamous cell carcinoma (OSCC) is the most common head and neck cancer and accounts for over 90% of cancers that develop in the mucosal epithelium of the oral cavity.^1–3^ Several carcinogenic risk factors, including tobacco and alcohol use, and viral agent infection (e.g. HPV) are associated with oral carcinogenesis.^4^ Unlike other head and cancer subsites such as the oropharynx, HPV accounts for only 2-5% of OSCC and the significance of HPV infection in OSCC is currently unknown.^5^ HPV-negative disease, mostly driven by tobacco and alcohol use accounts for a majority of OSCC and improving the prognosis of this subset is an area of unmet need.^4^ Despite advances in the understanding of OSCC biology over the past few decades, mortality outcomes have remained largely static and less than 50% of HPV-negative OSCC patients survive more than 5 years.^2^ Conventional treatment modalities such as surgery and cytotoxic chemotherapy have also experienced limited success and can result in severe morbidity,^6,7^ highlighting the need for alternative treatment strategies.

OSCC invasion and metastasis is poorly understood and accounts for a majority of cancer-associated deaths.^8^ More than 50% of patients experience cancer recurrence or develop metastases within 3 years of treatment.^8^ The OSCC leading edge, comprised of tumor cell layers at the border of the OSCC tumor, has been previously identified to have prognostic value in clinical grading and may mediate tumor invasion and metastasis.^9^ However, the active mechanisms at the invasive edge, and other important spatially-defined regions of carcinomas, are not fully understood.^10^ Previously, immunohistochemistry and in-situ hybridization efforts to study the leading edge have been limited to low-throughput analysis and have struggled to comprehensively characterize the OSCC tumor microenvironment.^9,11^

Recent advances in single-cell RNA sequencing (scRNA-seq) have enabled the exploration of intratumoral heterogeneity in head and neck squamous cell carcinoma (HNSCC).For instance, tumor cells expressing a partial epithelial mesenchymal transition (p-EMT) program were localized at the leading edge and demonstrated enhanced invasive potential.^12^ Conversely, tumor cells lacking p-EMT expression but expressing epithelial differentiation program markers were localized to the tumor core.^12^ However, scRNA-seq studies ultimately lack the spatial resolution that profoundly explores the biology of the tumor.^13^ Spatial Transcriptomics (ST) builds upon scRNA-seq by providing expression data while simultaneously preserving the 2D positional information of cells, providing an improved representation of transcriptional heterogeneity in the tumor microenvironment.^14^

Here, we leverage ST and single-cell RNAseq to unravel intratumoral transcriptional heterogeneity in OSCC by determining and characterizing the OSCC tumor core and leading edge in terms of their unique transcriptomic profile, cellular composition, and ligand-receptor interactions. Differences in regional expression may be explained by the presence of spatially unique cancer cell states. We also discovered that conserved transcriptional programs in the tumor core and leading edge have prognostic value not only in OSCC, but across multiple cancer types. Through predictive machine learning models, we also find that the leading edge regions may be conserved across cancer types, while the tumor core is more cancer specific. Furthermore, our work leverages RNA velocity inference to identify patterns of tumor differentiation within the tumor core and leading edge in relationship to drug response. Together, our results provide a novel outlook into the complex OSCC tumor landscape, and highlight the importance of further understanding the tumor core and leading edge in different cancer contexts for drug discovery.

## Results

### Conserved tumor core and leading edge transcriptional programs identified using ST

We performed ST on 12 surgically resected OSCC samples from 11 unique patients using the 10x Genomics Visium platform (Fig. 1a and Supplementary Table 1). In all, transcriptomes from 24,876 spots were sequenced to 43,648 post-normalization mean reads per spot. Data was normalized, corrected for batch effects across samples, dimensionality reduced, and louvain clustered for subsequent analysis. Hematoxylin and Eosin (H&E)-stained images from each tumor sample were examined and morphological regions were annotated by the study pathologist (M.H.) (Fig. 1b).

**Fig. 1:**
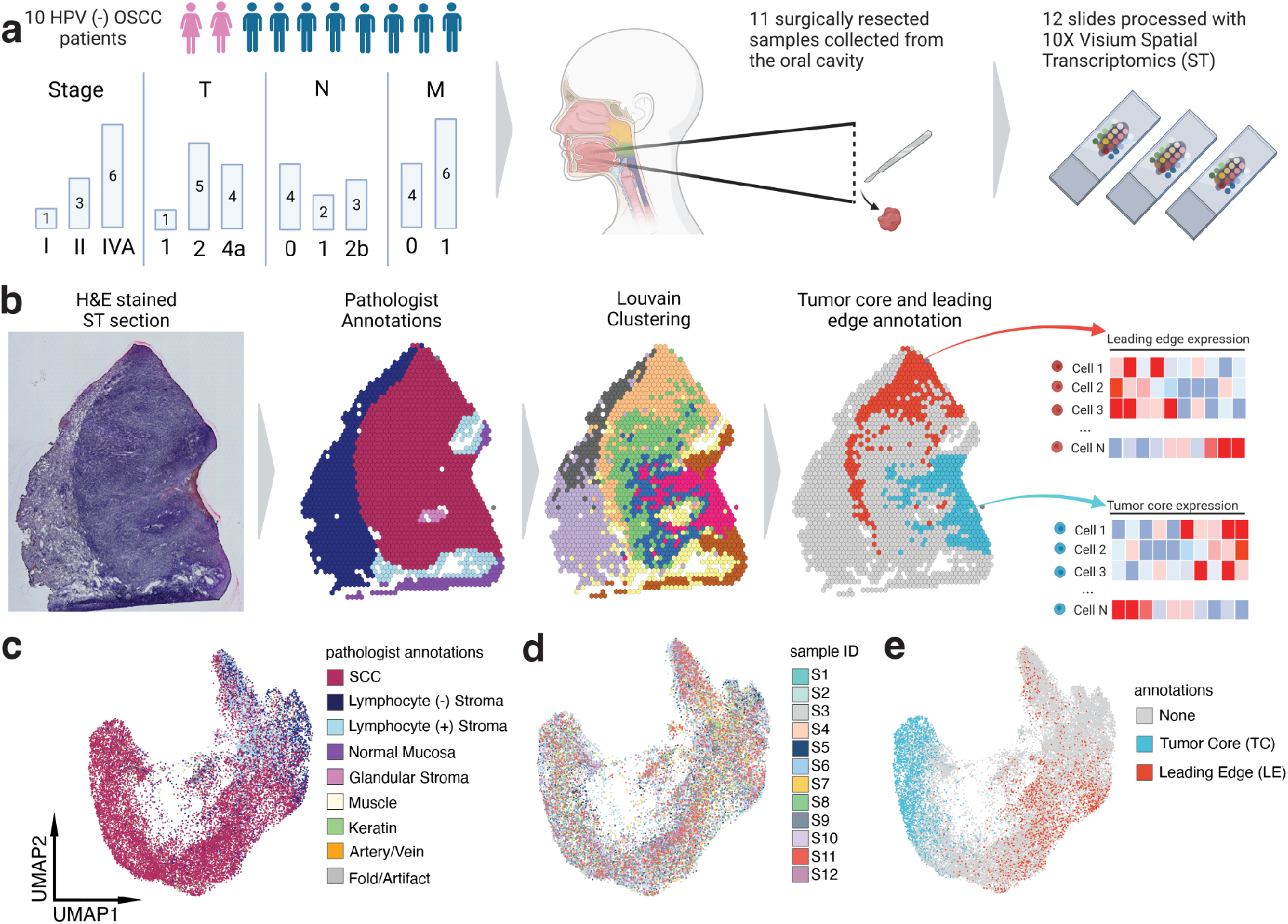
Experimental Overview of ST analysis on OSCC patient cohort and ST Analysis. **a**. Schematic representing patient clinical data and sample acquisition and processing strategy. **b**. A representative tissue section from an OSCC patient subjected to the ST pipeline; from left to right: a brightfield H&E image of the section, detailed pathologist annotations, spatially-naive Louvain clusters overlayed on pathologist annotations, TC and LE annotation strategy using TC and LE-expressed genes from the literature..**c-e**. UMAP projection of 24,876 spots aggregated from all 12 spatially-profiled samples colored based-on pathologist annotations, sample ID, and TC and LEannotations.

To determine the presence and location of the leading edge and tumor core in our OSCC samples, we scored pathologist-annotated tumor-associated spots in each unsupervised cluster for the expression of literature-identified genes localized to either spatial regions (Supplementary Table 2). Top scoring tumor-associated clusters were annotated as leading edge and tumor core, with 3043 tumor core spots and 3039 leading edge spots (Fig. 1b, Supplementary Table 1, and Extended Data Fig. 1a-i). Uniform manifold approximation and projections (UMAP) of ST spots revealed separation between tumor core and leading edge annotations, reflecting distinct transcriptional profiles between tumor core and leading edge regions within the OSCC tumor (Fig. 1c-e).

### The tumor core and leading edge are functionally heterogeneous components of the tumor microenvironment

Since tumor core and leading edge differences seemed visually conserved across patients in UMAP projections (Fig. 1b), we sought to determine whether the patterns of gene expression in the leading edge and tumor core are conserved across different patients. To do this, a correlation matrix was generated from the whole transcriptome gene expression profiles within the two spatial regions (Fig. 2a). A high degree of correlation was observed within the tumor core, and within the leading edge, across different patients. Interestingly, the correlation between the tumor core and leading edge expression programs within each patient was relatively low. Therefore, our tumor core and leading edge annotations comprise transcriptomically unique regions within tumors that are conserved across patients.

**Fig. 2:**
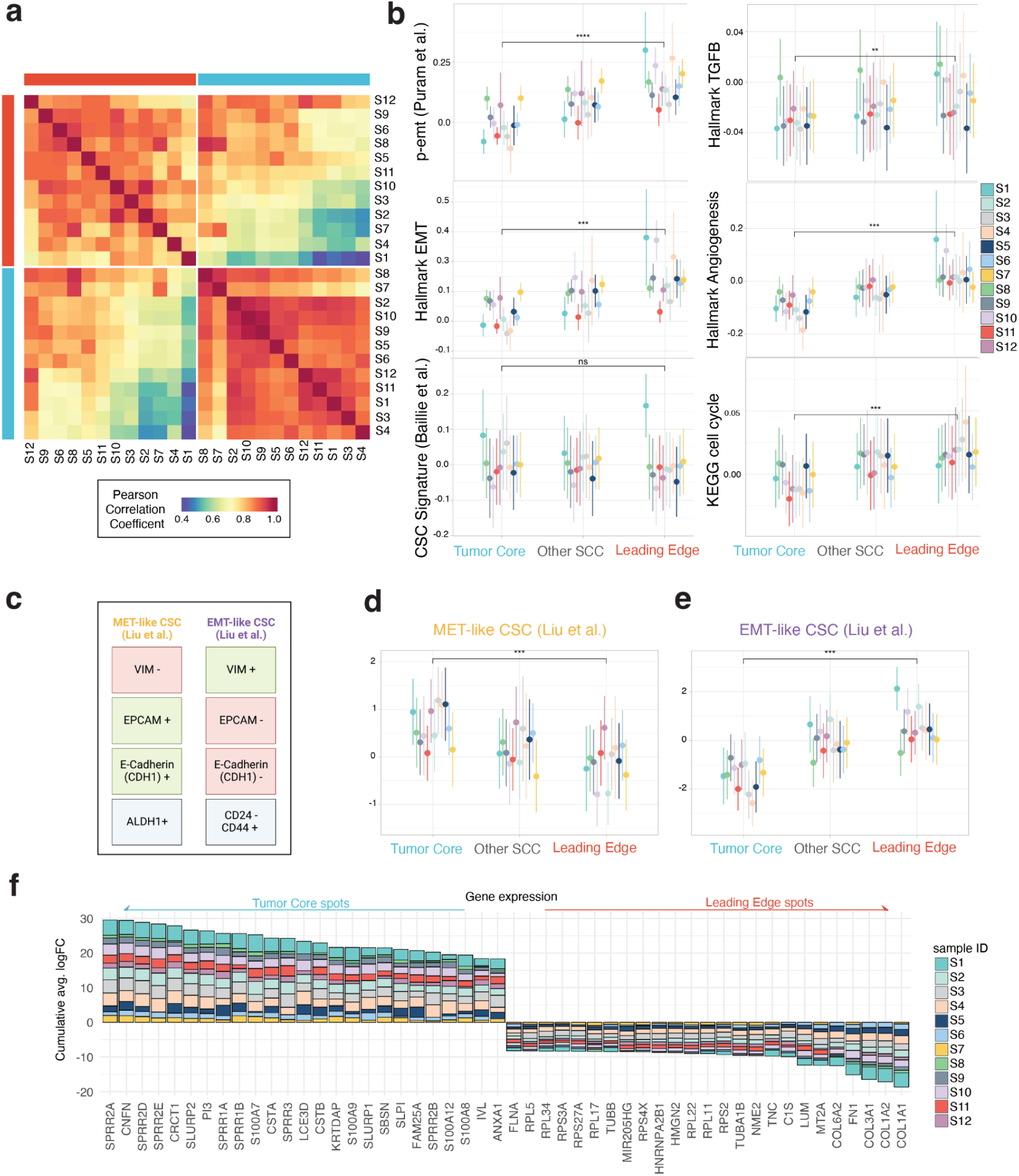
TC and LE are spatially unique regions in the OSCC microenvironment. **a**. Whole transcriptome correlation heatmap of TC and LE annotations across all spatially-profiled samples. Samples are ordered based on transcriptomic similarity. **b**. Comparative expression of six genesets across the TC, LE, and other squamous cell carcinoma (SCC) spots. All tested genesets had a significant p-value, except for the CSC signature. **c**. Schematic representation of mesenchymal (mCSC) and epithelial cancer stem cell (eCSC) markers characterized by Liu *et al*. **d-e**. Comparative expression of mCSCs and eCSC gene sets across the TC, LE, and other SCC spots. (For all gene set comparison plots, significance of differences between the tumor core and leading edge was determined through the use of a paired T test. Circles are representative of mean and lines are representative of inter-quartile range. All tested genesets had a significant p-value, *p < 0.05, **p < 0.01, ***p < 0.001, ****p < 0.0001.) **g**. Consensus plot displaying the cumulative average logFC for genes significantly (adj. p < 0.001) differentially expressed between the TC and LE across more than 9 samples.

We explored the functional differences between tumor core and leading edge spots. Querying cancer hallmark and key oncogenic pathway-associated genesets, we found that the tumor core and leading edge significantly differed in expression. Leading edge spots displayed higher expression of genes associated with cell cycle (p<0.001), TGF signaling (p<0.01), epithelial-mesenchymal transition (EMT) (p<0.001), and angiogenesis (p<0.001), in comparison to the tumor core (Fig. 2b). EMT scores in the leading edge were also expressed over a broad range (Fig. 2b), in alignment with the EMT continuum model.^10^ We additionally queried a published p-EMT geneset and observed localization of this program to leading edge spots (p<0.0001) (Fig. 2b).^12^ Ingenuity Pathway Analysis (IPA) predicted the activation of GP6, Actin Cytoskeleton, and Tumor Microenvironment canonical signaling pathways in the leading edge (Extended Data Fig. 2a). These characteristics might reflect the role of the OSCC leading edge in governing invasive behavior and metastasis. We then investigated the expression of cancer stem cells (CSC) within the tumor core and leading edge; CSCs are cancer cell populations that possess stem-cell like and malignant properties.^15^ No significant difference in expression of canonical OSCC CSC markers^16^ between the leading edge and tumor core was observed (p>0.05) (Fig. 2b). We hypothesized that unique CSC populations may be present in the tumor core and leading edge. We applied genesets associated with mesenchymal-like and epithelial-like CSCs (Fig 2c),^17^ and observed spatial localization to the tumor core (p<0.001) and leading edge (p<0.001), respectively (Fig 2. 2d,e). These findings corroborate the increased expression of the EMT signature in the leading edge. They also indicate that although tumor core and leading edge niches are both populated by CSCs, these CSCs are distinct in their biology.

We next explored regulatory differences between the tumor core and leading edge using single-cell regulatory network inference and clustering (SCENIC) to infer transcription factor (TF) activity. SCENIC analysis identified the upregulation of several proto-oncogenic TFs *CEBPA*, *JUNB*, and *KLF4* (ref.^18–20^), and tumor suppressor TFs *GRHL1* and *RCOR1* (ref.^21,22^) in the tumor core (Extended Data Fig. 2b and Supplementary Table 3). Interestingly, the upregulation of several TFs including invasion- and metastasis-regulating genes *RAD21* and *E2F4* (ref.^23,24^), and EMT regulatory genes *MAZ*, *ESSRA*, and *TWIST2* (ref.^25–27^) were observed in the leading edge (Extended data Fig. 2b and Supplementary Table 3).

Differential expression (DE) analysis between the tumor core and leading edge revealed 80 genes upregulated in the tumor core and 38 in the leading edge, across 10 or more samples (Fig. 2f and Supplementary Table 4). Upon comparison to a previous HNSCC scRNA-seq study,^12^ 37 epithelial differentiation and 4 p-EMT DEGs overlapped with our tumor core and leading edge DEGs, respectively. Top differentially expressed genes in the tumor core included genes involved in keratinization *SPRR2A*, *CNFN*, *SPRR2D*, and *SPRR2E*, while top genes in the leading edge included collagens (*COL1A1*, *COL1A2*, *COL3A1*, *COL6A2*) and fibronectin (*FN1*), highlighting the presence of a fibrovascular niche (Fig. 2f and Supplementary Table 4).

Four HNSCC molecular subtypes classified by patterns of gene expression and clinical outcomes have been identified and validated by the The Cancer Genome Atlas (TCGA).^28,29^ To determine if the unique expression profiles observed in the OSCC tumor core and leading edge could be attributed to the composition of HNSCC molecular subtypes, we integrated TCGA expression subtype signatures with our data. Spots within the tumor core and leading edge were scored for their correspondence to subtype expression signatures. We found that multiple molecular subtypes may be present within the same tumor microenvironment across different patients, with no consistent pattern of subtype composition (Extended Data Fig. 2c). Overall, the tumor core state was generally most enriched for the Basal subtype (p<0.0001) (Extended Data Fig. 2d), while the leading edge was generally most enriched for the mesenchymal subtype (p<0.0001) (Extended Data Fig. 2e). This suggests that differences between the tumor core and leading edge can not be explained by HNSCC molecular subtypes alone and that the absolute classification of patients into subtypes based on gene expression may not adequately represent their biology.

### Tumor core and leading edge gene signatures are conserved pan-cancer and are distinct in their prognostic impact

Since our annotated tumor core and leading edge regions are highly conserved across each of our OSCC samples, we wondered if these spatial regions were also present across other cancers as distinct molecular programs. We trained three machine learning (ML) probability based prediction models on our tumor core spots, leading edge spots, and all other remaining spots to generate a spatial region predictive model (Fig. 3a). We applied our prediction model to 13 samples across 9 different cancer types to characterize each spot in each sample (Fig. 3b). All 13 samples identified leading edge spots that were highly spatially segregated (Fig. 3b-e) (Extended data Fig. 3a). This is especially evident in cutaneous Squamous Cell Carcinoma (cSCC) samples and Hepatocellular Carcinoma (HCC) samples, where ML-predicted leading edge annotations corroborating those annotated by study authors (Fig. 3c,d).^30,31^ Only 2/13 sections identified tumor core spots that were spatially segregated (Fig, 3c,d) (Extended Data Fig. 3a), both of which corresponded to cSCC sections. Our results suggest for the first time that leading edge-associated expression states are conserved across multiple cancer contexts, while expression profiles associated with the tumor core are more tissue specific.

**Fig. 3:**
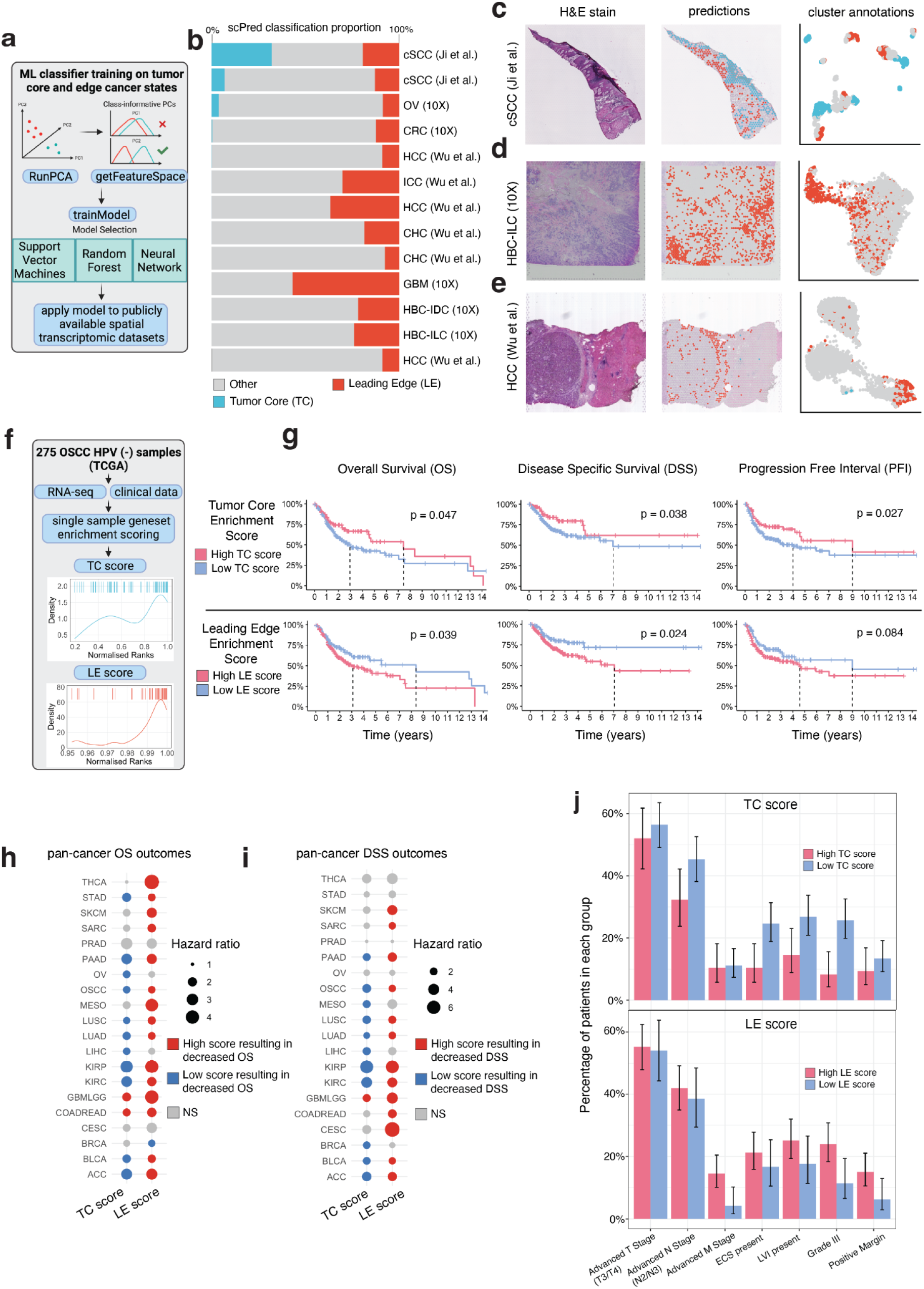
Pan-cancer conservation and survival associations of TC and LE transcriptomic signatures. **a**. Infographic describing the machine learning strategy used for the identification of TC and LE gene signatures in publicly-available spatially-profiled samples. **b**. Bar plot displaying the scPred classification score of each spatially distinct region across 13 different spatially profiled samples. The plot is ordered by the decreasing proportion of TC scores. **c-e**. H&E-stained tissue section (left), scPred projections on stained tissue (middle), and a UMAP colored by scPred classification (right) for cutaneous SCC (cSCC), Histiocytoid breast carcinoma Invasive Lobular Carcinoma HBC-ILC, and hepatocellular carcinoma (HCC) representative spatial transcriptomics testing datasets. **f**. Infographic describing TC and LE single-sample geneset scoring strategy for TCGA transcriptomic data. **g**. Kaplan-Meier visualizations of OS, DSS, and PFI end-points stratified by TC (upper panels) and LE (lower panels) gene set enrichment scores. P-values displayed were calculated using a log-rank test. **h**. OS and **i**. DSS pan-cancer outcomes for TC and LE geneset enrichment scores derived from 19 common cancer types from TCGA. P-values and hazard ratios displayed were calculated using a cox proportional-hazard regression. **j**. Bar plots showing the percentage of high and low TC (upper panel) and LE (lower panel) patients within relevant clinico-pathological covariates. Significance was determined using a hypergeometric test. Error bars represent standard error of mean (SEM).

To examine the prognostic significance of our leading edge and tumor core signatures, we incorporated samples from a bulk transcriptome dataset (TCGA) containing matched survival data. A total of 275 HPV-negative OSCC patients were selected for survival analysis and each sample was assigned an enrichment score based on the expression of genes differentially expressed in the tumor core and leading edge (see methods; Fig. 3f). Higher tumor core single-sample geneset scores were significantly associated with lower pathological staging (p<0.05) (Extended Data Fig. 4a), while leading edge scores did not significantly vary with pathological staging (p>0.05) (Extended Data Fig. 4b). Kaplan-Meier curves were generated to visualize the overall survival (OS), disease specific survival (DSS), and progression free interval (PFI) differences in relation to high or low expression of our tumor core and leading edge signatures (Fig. 3g). High expression of the leading edge signature was associated with worse OS and DSS in OSCC patients (p<0.05)(Fig. 3g). Conversely, high expression of the tumor core signature was associated with improved OS, DSS, and PFI (p<0.05) (Fig. 3g). Therefore, tumor core and leading edge signatures displayed starkly opposite survival outcomes. To validate our results, we performed the same comparison of high and low expression of tumor core and leading edge signatures in an independent cohort of 93 HPV-negative OSCC patients (GSE41613) and found the same trends across OS and DSS (Extended data Fig. 4c).

To determine if the prognostic impact of our leading edge and tumor core signatures was generalizable across different cancer types, we extended our analysis to 20 common solid tumors in TCGA. We found that a high leading edge score was consistently associated with worse OS and DSS across multiple cancers with the exception of breast cancer (BRCA) (Fig. 3h,i). In turn, a high tumor core signature was consistently associated with improved OS and DSS across multiple cancers with the exception of Glioblastoma and Lower Grade Glioma and (GBMLGG) and Colorectal Adenocarcinoma (COADREAD) (Fig. 3h,i). These findings suggest generalizability of our tumor core and leading edge expression programs across solid tumors and underline the presence of conserved spatial biological niches. However, the association of the tumor core and leading edge signatures with PFI was less consistent (Extended Data Fig. 4d). We next examined if our leading edge and tumor core signatures were associated with relevant clinical covariates in OSCC. High leading edge scores were associated with the presence of distant metastases (p<0.05), while low tumor core scores were associated with presence of extracapsular spread (p<0.05) and with higher tumor grade (p<0.05) (Fig. 3j).

### Distinct cancer cell states inhabit the tumor core and leading edge

To further determine the spatial architecture of tumor cells and other cellular subpopulations present in the tumor microenvironment, we integrated our ST data with a HNSCC scRNA-seq dataset containing malignant and immune cells (Extended Data Fig. 5a).^12^ CAF subtypes conserved in HNSCC were annotated using the marker genes *LRRC15* and *GBJ2* for ecm-MYCAFs, and *ADH1B* and *GPX3* for detox-iCAFs (Extended Data Fig. 5a).^32,33^ Spatial deconvolution analysis found that cancer, ecm-myCAF, detox-iCAF, dendritic, mast, macrophage, and cytotoxic CD8+ T cell types were represented in the tumor core and leading edge (Fig. 4a,b). The tumor core was primarily composed of cancer cells, with an extremely small proportion of CAFs in some samples (Fig. 4a,b). The leading edge was composed of cancer cells and ecm-myCAF cells, with a small proportion of detox-iCAFs, macrophages, and other immune cells (Fig. 4a,b). EPIC deconvolution of tumoral spots confirmed an increased proportion of CAF cells present in the leading edge (p<0.001) (Extended Data Fig. 5c). To identify the contribution of different cell populations in defining our tumor core and leading edge signatures, we projected the expression of our tumor core and leading edge DEGs with HNSCC scRNA-seq data. The tumor core signature was almost solely expressed in cancer cells (Extended Data Fig. 5d), while the leading edge signature was highly expressed in cancer cell and fibroblast cell populations (Extended Data Fig. 5b,c). These results suggest that the leading edge and tumor core microenvironments have different cellular compositions primarily defined by ecm-myCAFs and cancer cells. Consequently, we were curious to see if differences between the tumor core and leading edge were due to different proportions in ecm-myCAF presence or the potential existence of two distinct cancer cell states.

**Fig. 4:**
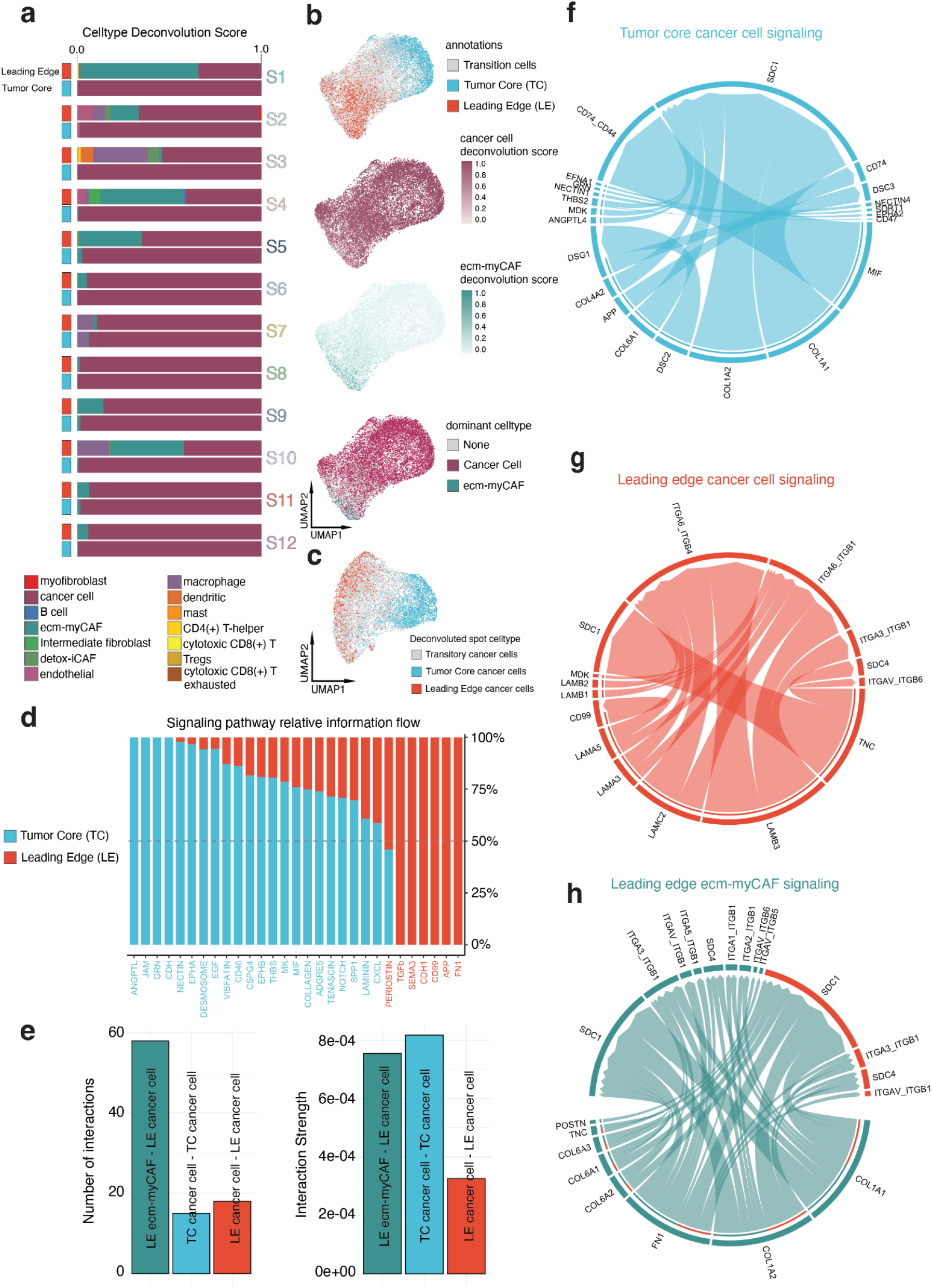
Integrated analysis of single-cell and spatial transcriptomics data reveals unique cellular composition, signaling pathways and cell-cell interactions in the TC and LE. **a**. Stacked bar plots visualizing the cumulative cell-type deconvolution scores within the TC and LEspots within each sample. Deconvolution scores were obtained through integration and label transfer of ST data with an OSCC single-cell RNAseq dataset. **b**. UMAP of tumor microenvironment-associated spots aggregated from all ST profiled samples colored by (from top to bottom) TC and LE annotations, cancer cell deconvolution score, ecm-myCAF deconvolution score, and dominant deconvoluted cell-type. **c**. UMAP of spatially deconvolved cancer cell only spots, colored by TC and LE annotations. **d**. Stacked bar plot visualizing significantly enriched signaling pathways by overall information flow and their activities across TC and LE spots. Pathways are colored by the dominant within the TC or LE. **e**. Bar plot visualizing number and cumulative strength of inferred ligand-receptor interactions for spatially deconvolved LE ecm-myCAFs with LE cancer cells, TC cancer cells with TC cancer cells, and LE cancer cells with LE cancer cells. Interaction strength represents the probability of ligand-receptor interaction. Circos plots describing spatially deconvolved ligand-receptor pairs involved in cell-cell interactions between **f**. cancer cells in the TC, **g**. cancer cells in the LE, and **h**. ecm-myCAF signaling and cancer cells in the LE. The width of connecting bands represents the strength of ligand-receptor interaction.

To explore this possibility, we stringently characterized cancer cells across both the tumor core and leading edge as having a deconvolution score of > 0.9. Seven samples were identified to have both spatially deconvolved tumor core and leading edge cancer cells based on the applied cutoff with high confidence. Following batch effect correction and dimensionality reduction of our classified cells, UMAP projections showed separation between spatially deconvolved tumor core and leading edge cancer cells, reflecting distinct transcriptional profiles (Fig. 4c). We found the expression profiles within tumor core cancer cells (TCC), and within leading edge cancer cells (LEC), were highly similar across patients (Extended Data Fig. 5e). However, a lower degree of similarity was seen between comparisons of TCC and LEC expression profiles across patients (Extended Data Fig. 5e). Expression profiles of ecm-myCAFs found in the leading edge also differed from LEC and TCC (Extended Data Fig. 5e). Subsequent differential expression analysis found 115 genes upregulated in TCCs and 99 in LECs across all patients featuring both LECs and TCCs (Extended Data Fig. 5f and Supplementary Table 5). Taken together, these findings suggest the presence of transcriptionally unique cancer cell states within the tumor core and the leading edge may contribute to their distinct biologies.

### The tumor core and leading edge display distinct ligand-receptor interactions

Given the considerably diverse spatial architecture of the OSCC tumor core and leading edge, we sought to elucidate the nature and role of intracellular and extracellular signaling pathways in the tumor core and leading edge. We performed cell-cell communication analysis with the CellChat package to derive quantitative inferences of intercellular communication networks (Supplementary Table 6). After quantifying and visualizing global patterns of cell-cell communication within OSCC samples, the greatest inferred strength of outgoing information was seen in the leading edge, and incoming information in the tumor core (Extended Data Fig. 6a). ANGPTL, JAM, GRN, and CDH signaling pathways were exclusively seen in the tumor core, and TGFB, SEMA3, CDH1, CD99, APP, and FN1 in the leading edge (Fig. 4d). Several ECM remodeling pathways including Collagen, Tenascin, and Laminin were also expressed in both the tumor core and leading edge (Fig. 4d).

We next examined cell-cell communication mechanisms within our spatially deconvolved data. Leading edge ecm-myCAFs exhibited diverse signaling with many more interactions to LEC compared to TCC-TCC and LEC-LEC signaling. Moreover, leading edge ecm-myCAF-LEC interaction strength was similar to TCC-TCC interaction strength and greatly exceeded the interaction strength for LEC-LEC signaling (Fig. 4e). The relatively greater number of ecm-myCAF-LEC interactions and strength relative to LEC-LEC signaling highlights the critical role of ecm-myCAFs in facilitating biological features of OSCC tumors. Outgoing signaling patterns exclusive to LEC-LEC signaling, relative to TCC-TCC signaling, included Laminin, Tenascin, and CXCL pathways (Extended Data Fig. 6b), reinforcing the role of these pathways in facilitating cancer invasion and metastasis.

We then explored specific ligand-receptor pairs representative of TCC-TCC and LEC-LEC signaling. We found that TCC could signal to TCC by adhesive ligand-receptor pairs COL1A2/COL1A1-SDC1 and inflammatory ligand-receptor pairs MIF-CD74_CD44 (Fig. 4f and Supplementary Table 6). Interestingly, the MIF-CD74 ligand-receptor pair has been previously identified to initiate oncogenic signaling pathways.^34,35^ Similarly, LEC could contact LEC through adhesive ligand-receptor pairs LAMB3-ITGA6_ITGB4 and LAMB3-ITGA6_ITGB1 (Fig. 4g and Supplementary Table 6). Strikingly, there were no ligand-receptor pairs that overlapped between TCC-TCC and LEC-LEC signaling. Due to the high number of leading edge ecm-myCAF-LEC interactions, we also analyzed the signaling pathways between LECs and leading edge ecm-myCAF cells. Our analysis predicted prominent signaling mediated by ecm-myCAF-LEC ligand-receptor pair interactions, including adhesive COL1A1/COL1A2-SDC1 and FN1-SDC1 ligand-receptor interactions (Fig. 4h and Supplementary Table 6). Taken together, our findings suggest that the OSCC tumor core and leading edge have distinct patterns of cell-cell communication that extend beyond their unique cellular compositions and further highlight their unique biologies.

### Differentiation trajectories revealed by analyzing RNA splicing dynamics between the tumor core and leading edge

RNA velocity can be used to predict the short-term future development of individual cells using the ratio of spliced and unspliced mRNA counts.^36^ Aggregating results together helps reveal the developmental trajectories of cancer cells, and similarly identify putative driver genes responsible for the transition.^37^ We utilized RNA velocity to characterize the development of cancer cells present in the tumor core and leading edge. Among spatially deconvolved cancer cells aggregated across all samples, we observed a differentiation hierarchy originating from TCC extending towards LEC (Fig. 5a). Similar patterns of directional flow were also observed at the individual patient level (Fig. 5b). Several genes displayed dynamic splicing behavior that drove the core to edge cancer cell differentiation hierarchy. Among the top putative driver genes were the *CSTA* and *MT2A* genes, the splicing of which was higher in the tumor core and leading edge, respectively (Fig. 5c,d and Supplementary Table 7). *CSTA* has been identified as a tumor suppressor gene involved in mesenchymal-epithelial-transition (MET), while *MT2A* is a key regulator of EMT.^38,39^ These findings suggest that the tumor core is predicted to undergo increased differentiation retaining an epithelial phenotype, while the leading edge becomes increasingly mesenchymal. Other differentially spliced driver genes in the tumor core and leading edge included proto-oncogenes and tumor suppressor genes (Fig. 5c, and Extended Data Fig. 7a and Supplementary Table 7). Spots aggregated from the entire OSCC tumor (including non-malignant cells) also exhibited a similar differentiation trajectory, with differentiation occurring from the tumor core into the leading edge (Fig. 5e).

**Fig. 5:**
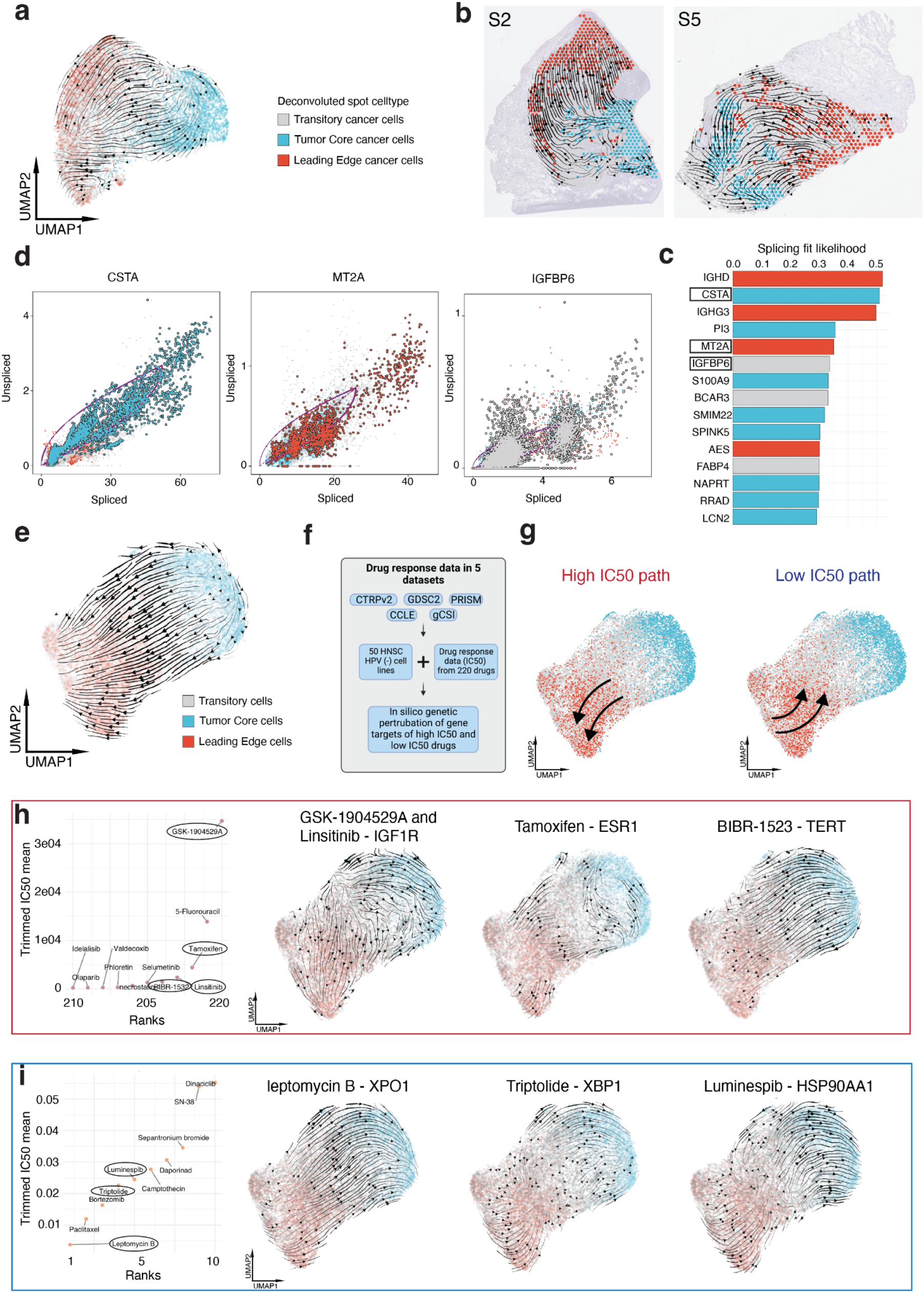
RNA dynamics analyses reveal differential developmental trajectories and therapeutic vulnerabilities in the TC and LE. **a**. UMAP of spatially deconvolved cancer cell spots, with overlaid RNA velocity streams, colored based on TC and LE annotations. **b**. Representative spatially profiled samples (samples 2 and 5) overlayed with RNA velocity streams and colored by TC and LE cancer cell annotations. **c**. Bar plot visualizing top differentially spliced genes (genes with dynamic splicing behavior) within TC and LE regions. Bars are colored by whether a higher proportion of the gene exists in its spliced form in the TC or LE. **d**. Phase portraits showing the ratio of spliced and unspliced RNA for top differentially spliced genes, purple lines depict predicted splicing steady state. **e**. RNA velocity analysis of tumor-associated spots colored by TC and LE annotations aggregated across all spatially profiled samples. **f**. Infographic describing the strategy used to identify effective (low IC50) and ineffective (high IC50) anticancer drugs. **g**. UMAP plot from e. overlaid with summary response curves based on vector field perturbations observed in h and i. Rank ordered dotplots and UMAPs showing the resultant vector field following *in silico* perturbations of three targets of **h**. high (ineffective) IC50 anticancer drugs, and **i**. low IC50 anticancer drugs..

### Deriving therapeutic targets for OSCC

OSCC is characterized by its high disease burden and mortality, despite improvements in diagnosis and treatment modalities.^40^ Existing treatments such as surgery, chemotherapy and radiotherapy are effective only in a minority of patients and OSCC recurrence remains a leading cause of death.^40^ Furthermore, current treatments may induce significant morbidity and reduction in the quality-of-life (QoL) related to non-specific cell death, including local defects, speech and swallowing dysfunction, and other toxic side-effects.^40,41^ Targeted therapies have recently gained attention for their ability to mitigate host toxicity while improving survival rates and QoL.^40^ However, few targeted therapies are currently used in OSCC management.^42^ Therefore, we applied an in-silico based perturbation approach to identify spatially regulated families of RNA velocity drug responses that may predict therapeutic success in the OSCC tumor (Fig. 5f). Dynamo is an *in-silico* technique that is capable of accurately predicting cell-fate transition following genetic perturbation based on learned splicing vector fields.^43^ PharmacoDB drug-response data^44^ for 220 drugs across at least 25 HPV negative HNSCC cell lines was rank-ordered based on trimmed IC50 means (Supplementary Table 8). Among ineffective drugs (high IC50), dynamo based *in-silico* perturbations displayed consistent patterns of directional flow extending from intermediary regions to the leading edge, similar to the baseline vector field (Fig. 5g,h and Extended Data Fig. 7d). Conversely, dynamo based *in-silico* perturbations of effective drugs (low IC50) had consistent patterns of directional flow extending from the leading edge to intermediary regions (Fig. 5g,i and Extended Data Fig. 7e). Interestingly, dynamo based *in-silico* perturbations of common immunotherapy targets (anti-PD-1, anti-ctla-4) displayed similar results to effective drugs (low IC50) with patterns of vector flow extending from the leading edge to intermediary regions (Extended Data Fig. 7b,c).

## Discussion

The spatial organization of OSCC tumor microenvironment is poorly understood and there are limited insights on how the tumor core and leading edge affect OSCC biology. Here, we leverage spatial transcriptomics profiling of 12 OSCC tissue samples to extensively characterize the tumor core and leading edge. Our novel analysis unraveled the complex and unique spatial architecture, cellular composition, ligand-receptor interactions, prognostic impact and potential druggability of gene expression programs within the OSCC tumor core and leading edge– insights which are applicable pan-cancer.

Previous HNSCC scRNA-seq studies have revealed the importance of several gene expression programs and their spatial location within the tumor.^12^ In this study, we find that the OSCC tumor core is characterized by a keratinized and differentiated state, while the leading edge confers several invasive and metastatic properties. In agreement with Liu et al, we found that epithelial-like CSCs were localized to the tumor core while mesenchymal-like CSCs to the tumor leading edge.^17^ We also find the differences in gene expression between the tumor core and leading edge are not governed by HNSCC molecular subtype compositions, and that multiple subtypes may be reprepresented within a single tumor sample. Rather, transcriptomic differences may be attributed to the unique compositions of cancer cells and fibroblasts, among other factors. A recent study has identified the existence of conserved cancer cell states across different cancers due to unique patterns of gene module expression.^45^ Another scRNA-seq study in OSCC has also alluded to the existence of several unique tumor cell populations.^46^ Therefore, we propose that differences in the tumor core and leading edge are driven by the existence of spatially unique cancer cell states. Through velocity analysis, we find that these spatially unique cancer states are governed by a differentiation hierarchy composed of progenitor cancer cell states from the tumor core developing into more specialized cancer cell states of the leading edge. Taken together with the distinct CSC contexture in the tumor core (epithelial-like CSCs) versus the leading edge (mesenchymal-like CSCs), we believe that cancer cells from the core transition into leading edge cells by gradually acquiring a more aggressive EMT-like phenotype that helps in invasion and dissemination. Although multiple previous studies have described the presence of transcriptionally unique cancer cell states,^12,45–47^ our research is the first to show that these states may also be spatially governed.

Our ML model served as a useful tool to explore the tumor core and leading edge in different cancer contexts. We found that the leading edge may exist across several different cancers and can be annotated with considerable accuracy, while the programs present in the tumor core may be reducible to tissues with similar origins. To our knowledge, this phenomenon has not been previously described. These findings may direct opportunities to better understand the conserved features in cancer invasion and metastasis, and guide subsequent treatment efforts targeted at the leading edge in a pan-cancer scope.

Differentially expressed genes in the OSCC tumor core and leading edge were found to be valuable for prognostication. We found that high leading edge signature scores and low tumor core scores were associated with worse patient outcomes. The tumor core may also confer its protective effect by altering tumor differentiation. To the best of our knowledge, our study is the first to describe that gene expression programs within the tumor core have a protective effect on survival. We also find that the prognostic ability of our leading edge and tumor core gene signatures may be generalizable across different cancer contexts. We hypothesize that the leading edge signature may be particularly relevant across multiple cancers, while the tumor core signature may be specific to epithelial keratinizing cancers.

Finally, we illustrate a novel application of RNA velocity to predict drug success. Through an *in-silico* based perturbation approach, we illustrate how different drugs with varying levels of effectiveness, may induce unique spatial drug responses. Effective drugs (low IC50 levels) consistently demonstrated RNA velocity patterns that opposed canonical differentiation patterns in the edge, and extended from the leading edge to the tumor core. Conversely, ineffective drugs (high IC50 levels) generally agreed with canonical differentiation patterns in the edge that extended from the tumor core to the leading edge. Importantly, the results of our spatial *in-silico* perturbation analysis show that drug-responses occur mainly within the leading edge, highlighting the tumor leading edge as a credible drug target. Our findings ultimately provide an exciting *in-silico* based approach which may be able to identify novel anticancer drugs.

Together, these findings suggest that spatially unique cancer cells exist within the OSCC tumor and play a role in shaping patient biology, outcomes and drug response. We propose that the tumor core functions as a reservoir of cancer cells in a precursor-like state while the leading edge plays a larger role in shaping cancer outcomes. The progenitor programs in the tumor core may maintain a level of benefit or survival advantage, while pathogenic mechanisms in the leading edge should be the target of therapies. The potential role of clonality in incurring the protective feature of the tumor core is unclear and remains to be explored in future studies. Our work represents the first effort to characterize and uncover unique spatial patterns of gene expression between the OSCC tumor core and leading edge, and the first application of ST to OSCC. The spatiotemporal mechanistic insights gained from our study will help direct the development of improved targeted therapies for OSCC and beyond.

## Methods

### Sample collection and annotation

Fresh-frozen OSCC tissue was obtained from the Alberta Cancer Research Biobank under the approval of the Health Research Ethics Board of Alberta – Cancer Committee (reference number: HREBA.CC-16-0644). The BioBank OSCC tumor samples were collected at the time of the surgery, embedded in an optimal cutting temperature (OCT) compound, and stored frozen at −80C until retrieval for the study. The study samples were sectioned in a cryostat at 10 um, with a first section used for ST and the next consecutive level stained with hematoxylin and eosin (H&E). The H&E sections were used to confirm the presence of an edge of OSCC and to annotate the tissues by the study pathologist (M.H). Regions were labeled as: oral squamous cell carcinoma tumor, lymphocyte-positive stroma, lymphocyte-negative stroma, normal mucosa, glandular stroma, muscle, keratin, artery/vein, and artifact. Pathologist annotations were exported from the loupe browser 6 and imported into Seurat for further analysis.

### Spatial transcriptomic profiling

Tissue optimization was performed to determine the optimal permeabilization time for OSCC tissue for the downstream gene expression protocol. Spatial transcriptomics was performed on OSCC cryosections using the Visium platform according to the manufacturer’s protocol (10x Genomics, Pleasanton, CA, USA). Briefly, OCT-embedded 10 micrometer-thick cryosections of OSCC samples were placed on the Visium spatial slide. Sections were enzymatically permeabilized for 24 min. cDNA was obtained from mRNA bound to capture oligos printed on the slide. cDNA quantification was performed using the Agilent Bioanalyzer High Sensitivity Kit on an Agilent Bioanalyzer 2100 (Agilent Technologies, CA, USA). cDNA libraries were sequenced on an Illumina NovaSeq 6000 sequencer using the SP flowcell (200 cycles) at the Centre for Health Genomics and Informatics (CHGI, University of Calgary, Alberta, Canada). Sequencing reads were aligned using the 10s Genomics Space Ranger 1.3.1 pipeline to the standard GRCh38 reference genome. 12 samples passing alignment QC were aggregated together using the 10x Genomics Space Ranger aggr function to normalize for read depth between samples. Aggregated samples (n = 12) recovered a total number of 24,876 spots containing tissue sequenced to 43,648 post-normalization mean reads per spot.

### Spatial transcriptomic data pre-processing

The aggregated HDF5 matrix was imported into R and split by sample. Feature-barcode matrices for each sample were imported into the R package Seurat for normalization, quality control, batch effect correction, dimensionality reduction, and Louvain clustering.^48^ Spots expressing less than 200 features were excluded from downstream analysis. Sample level normalization was performed using the SCTransform function in Seurat. Batch effect correction, integration, and dimensionality were performed using the introduction to scRNA-seq integration Seurat vignette (https://satijalab.org/seurat/articles/integration_introduction.html) with no deviations.

### Tumor core and leading-edge annotation

A literature search was conducted to identify genes expressed in the core^11,12,49^ and periphery of OSCC tumors.^9,11,12,50–52^ 15 leading edge genes and 7 tumor core genes were identified across multiple studies and scored in each spot using the Seurat addmodulescore function (Supplementary Table 2). Louvain clusters containing SCC with high leading edge geneset scores localized to the tumor border were annotated as the tumor leading edge. Louvain clusters containing SCC with high tumor core geneset scores localized to the center of the tumor were annotated as the tumor core. UMAP visualization of geneset scores was performed with Seurat and a first quartile minimum cutoff was used to reduce geneset noise. Annotated spots in sample 2 were visualized using the BayesSpace package for an infographic.^53^

### Differential expression analysis, consensus plotting, and correlation heatmaps

Differentially expressed genes were identified using a Wilcoxon rank-sum test in Seurat with a logFC > 0.25 and adjusted P value < 0.001. Stacked bar plots showing cumulative expression logFC for each gene across all samples were generated using an adaptation of the constructConsensus function.^54^ Consensus plots display top 25 genes differentially expressed in more than 9 samples (>9/12 samples). Differentially expressed genes for each sample were imported into Ingenuity Pathway Analysis for pathway enrichment analysis. Ingenuity pathway analysis exports were imported into the multienrichjam R package (https://github.com/jmw86069/multienrichjam). The multienrichjam mem_enrichment_heatmap function was modified to create pathway enrichment plots across samples. The whole transcriptome average expression was Pearson correlated using the cor function in the R stats package. Correlation values were plotted using the Heatmap function in the ComplexHeatmap R package.^55^

### Statistical approach for comparing scores across sample groups

Each gene signature was scored in each spot using the Seurat function addmodulescore. To test for differences in the module scores calculated between the tumor core and leading edge, a paired T test was conducted using the ggpubr function stat_compare_means. To perform the scoring of epithelial CSC and mesenchymal CSC signatures, an expression module was programmed. CSC gene characteristics were identified from literature according to their predicted upregulation or downregulation.^17^ The scoring contribution of genes predicted to be upregulated and downregulated were then cumulatively added to derive module scores.

### Gene regulatory network inference

SCENIC was used to infer the upstream regulatory activation of transcription factors. Tumor core and leading edge spots were subsetted from the Seurat object and processed in accordance with the SCENIC vignette with the standard hg19 RcisTarget reference (https://github.com/aertslab/SCENIC).^56^ A stacked bar plot showing the cumulative logFC of AUCell scores for each TF across all samples was generated using an adaptation of the constructConsensus function with an adjusted p value < 0.05.

### TCGA analysis and geneset scoring

Patient metadata, survival data, and bulk RNA sequencing data were downloaded for all samples in The National Cancer Institute’s Cancer Genome Atlas using the UCSCXenaTools R package.^57,58^ 275 OSCC HPV negative oral cancer patients were identified based on patient metadata. Gene set enrichment analysis was performed on genes differentially expressed in the tumor core and leading edge using the singscore R package with default parameters.^59^ In the tumor core, geneset enrichment scores greater than the 0.65 quantile were labeled as high and geneset scores less than the 0.65 quantile were labeled as low. In the tumor leading edge, geneset enrichment scores greater than the 0.35 quantile were labeled as high and geneset scores less than the 0.35 quantile were labeled as low. Kaplan-meier survival plots were generated using the survfit function in the survival R package and plotted using the ggsurvplot function in the survminer R package (https://github.com/kassambara/survminer). Hazard ratios and p-values were obtained from cox proportional hazard testing using the coxph function in the survival R package.

When comparing survival across multiple cancer types, the optimal quantile cut point for each geneset enrichment score in each cancer was determined using the surv_cutpoint function in the survminer R package. Optimized hazard ratios and p-values were subsequently calculated using the coxph function in the survival R package with default parameters.

To further evaluate our tumor core and leading edge genesets across multiple relevant clinical features, we adapted a previous approach used by Puram et al. and performed a hypergeometric test to compare the percentage of patients with high and low enrichment scores for both the tumor core and leading edge.^12^

### TCGA validation

To validate our findings in an external database, data from GSE41613 containing 93 HPV negative OSCC patients was downloaded and imported into R.^60^ Data was processed similarly to TCGA data; genesets were scored with singscore, stratified into high and low tumor core and leading edge scores using an optimized cutpoint, and had their optimized hazard ratios and p-values calculated.

### TCGA subtyping

TCGA subtype data for 279 OSCC samples was downloaded using the PanCancerAtlas_subtypes function from the TCGAbiolinks R package.^61^ Bulk RNA sequencing data and subtype data from the TCGAbiolinks package was added to a Seurat object using the CreateSeuratObject function.^61^ Deconvolution of spatial transcriptomic spots into HNSC subtypes was performed using the Seurat vignette for mapping and annotating query datasets with no modification (https://satijalab.org/seurat/articles/integration_mapping.html). The subtyped data was held as a reference dataset and the spatial transcriptomics data submitted as a query dataset.

TCGA subtype data was scored using the singscore R package for tumor core and leading edge genesets. Scores across HNSC subtypes were compared using a Kruskal-Wallis test implemented through the ggpubr function stat_compare_means.^59^

### Machine learning model for tumor core and leading edge

To assess whether tumor core and edge identified in OSCC were conserved in other solid tumors, we trained three machine-learning probability-based prediction models (Support Vector Machines with Radial Basis Function Kernel, Model Averaged Neural Network, and Random Forest) using scPred.^62^ Briefly, feature selection was performed by shortlisting top 50 class-informative PCs distinguishing spot-level variance between ‘tumor core’, ‘leading edge’, and ‘other’. Prediction models were trained using the caret R package. Training probabilities for spatial tumor states in the OSCC dataset were evaluated using get_scpred() and visualized using plot_probabilities(). 2 combined hepatocellular cholangiocarcinoma (CHC) (Wu et al.), 3 hepatocellular carcinoma (HCC) (Wu et al.), 1 intrahepatic cholangiocarcinoma (ICC) (Wu et al.), 2 cutaneous squamous cell carcinoma (cSCC) (Ji et al.), 1 glioblastoma (GBM) (10X genomics), 1 human breast cancer Invasive ductal carcinoma (HBC-IDC) (10X genomics), 1 human breast cancer Invasive lobular carcinoma (HBC-ILC) (10X genomics), 1 colorectal cancer (CRC) (10x genomics), and 1 ovarian cancer (OV) (10x Genomics) were classified and Harmony-integrated using scPredict() with default probability threshold.^30,31,63^ scPred-generated annotations were imported into Seurat and overlaid on top of histologic image using SpatialDimPlot(). The trained model used for classification is publicly available via our Figshare portal (https://doi.org/10.6084/m9.figshare.20304456).

### Cell type deconvolution

Single-cell HNSCC data was downloaded from GSE103322 and imported into R using the CreateSeurat object function in Seurat.^12^ In brief, the single-cell data was normalized using SCTransform, dimensionality reduced, and Louvain clustered. Cell types were assigned to Louvain clusters based on marker genes described in the included in the datasets original paper. CAF subtypes conserved in HNSC were annotated using the marker genes *LRRC15* and *GBJ2* for ecm-MYCAFs and *ADH1B* and *GPX3* for detox-iCAFs.^32,33^ Deconvolution of spatial transcriptomic spots into single-cell cell types was performed using the Seurat vignette for mapping and annotating query datasets with no modification (https://satijalab.org/seurat/articles/integration_mapping.html). The single cell dataset was held as a reference while the spatial transcriptomics data was submitted as a query dataset.

Ecm-myCAF dominant spots were identified as spots with an ecm-myCAF score of greater than 0.5 (with a max score of 1.0). Cancer cell dominant spots were defined more strictly as spots with a cancer cell score of greater than 0.9 (with a max score of 1.0). Cancer cell dominant spots were subsetted, corrected for batch effects, and dimensionality reduced as previously described. Celltype deconvolution was validated using the “epic” deconvolution method through the deconvolute function in the immunedeconv R package with default settings.^64,65^

### Inferring cell communication networks

The CellChat package was used to infer cell-cell interaction networks from a Seurat object containing deconvoluted spots using the createCellChat function.^66^ Filtered Circos plots were generated using the netVisual_chord_gene function in CellChat to visualize ligand-receptor pairs. Cellchat was also used to compare the overall information flow of the tumor core and leading edge in different signaling families using the rankNet function in comparison mode.^66^

### Constructing cellular trajectories using RNA velocity

To generate spliced and unspliced assays used to infer RNA velocity, the command line interface tool in velocyto was employed (https://velocyto.org).^67^ For each sample, barcodes and bam files corresponding to pathologist annotated cancerous regions were supplied to the velocyto “run” command. The velocyto “run” command was provided with a .gtf gene annotation file that was created using the cellranger mkref function applied to the standard GRCh38 reference genome. Output loom files were combined and imported into the scVelo python package for further analysis, dimensionality reduction coordinates and sample metadata for scvelo analyses were imported from Seurat.^37^ Multiple scVelo objects were processed in parallel identically. Spot-level velocity was derived from the scVelo dynamical model which identifies cellular trajectories based on the ratio of spliced mRNA to un-spliced pre-mRNAs. First and second order velocity vector moments were calculated using scv.pp.moments(data,n_pcs=10, n_neighbors=30). Dynamical velocity was calculated using the scv.tl.recover_dynamics function and the scv.t.velocity functions. Differential splicing for each gene was calculated using the scv.tl.rank_velocity_genes function which utilizes t-tests to compare multiple grouping variables and identify group-specific rank ordered gene lists. Data from scVelo was subsequently imported into the Dynamo python package for vector field learning and cell fate trajectory inference with no deviation from their standard vignette.^43^ Vector field topologies were learnt and visualized using the dyn.pl.topography function with default parameters and subsequently animated using the dyn.mv.animate_fates function.

### Drug response and in-silico perturbation of RNA velocity trajectories

Cell line drug response data was downloaded from the CTRPv2, GDSC2, PRISM, CCLE, and gCSI databases using the R package PharmacoGx.^68^ 50 HPV negative HNSCC cell lines were identified across all 5 drug datasets. 220 drugs were identified to have IC50 values available in at least 25 HPV negative HNSCC cell lines. IC50 values for each cell line were averaged across datasets for datasets with overlapping cell lines. The IC50 value for each drug was calculated by taking the 10% trimmed mean of the IC50 values observed across all cell lines to control for drug response outliers. Gene targets of drugs with the top 5 highest and bottom 5 lowest IC50 (only drugs with a genetic target) were identified in *silico* perturbation analysis with Dyanmo. In brief, genes were perturbed *in silico* using the Dynamo function dyn.pd.pertrubation with a Jv scaling factor of −1000.

### Spatial Transcriptomics Atlas

A web portal was created to enable exploration of all 12 spatial samples. This portal was built using RShiny, shinyLP, and shinythemes R packages. The portal is available for public access at http://www.pboselab.ca/spatial_OSCC/.

## Acknowledgements

This study was funded by the PRecision Oral Biology (PROBE) Grant from the Ohson Research Initiative and Charbonneau Cancer Institute to PB. We thank the Centre for Health Genomics and Informatics (CHGI) for providing the sequencing infrastructure to spatially profile our RNAseq libraries. We would like to thank the University of Calgary High Performance Computing Cluster (ARC) for providing data analysis infrastructure. We thank Danielle Simonot for her help with tissue retrieval from the Alberta Cancer Research Biobank (ACRB). We thank Christina Yang for her help with cryosectioning and slide preparation and H&E staining of profiled samples. We thank Dr. Mayi Arcellana-Panlilio and Dr. Guido van Marle for their extremely kind mentorship and productive feedback. We would like to thank Arzina Jaffer and Keerthana Chockalingam for help with setting up bioinformatics tools and bioinformatics consultation. We would also like to thank the patients and their families for consenting to provide tissue for this study.

## Data availability

Raw and SpaceRanger processed spatial transcriptomics files will be available upon publication at the National Center of Biotechnology Information’s Gene Expression Omnibus (GEOXXXX). Processed Seurat objects, loom files generated by velocyto, and the scPred prediction model will be available for reanalysis at figshare upon publication. Geneset IDs M5930, M7963, M5944, and M2642 were collected from the Molecular Signatures Database v7.5.1.^69^ P-emt and cancer stem cell genesets were collected from prior studies.^12,16,17^ Bulk RNA-sequencing data and associated clinical data used for survival analysis were downloaded from the National Cancer Institute’s The Cancer Genome Atlas.^58^ A validation genomic survival dataset was downloaded from GSE41613.^60^ Datasets used for scored analysis are publicly available and were downloaded from http://lifeome.net/supp/livercancer-st/data.htm (liver cancer),^30^ GEO accession number GSE144240,^31^ and 10X datasets downloaded from https://www.10xgenomics.com/resources/datasets (OV, CRC, HBC, GBM). Drug response data was downloaded from Pharmacodb.^44^ Spatial datasets are also available for public access at our companion portal http://www.pboselab.ca/spatial_OSCC/.

## Code availability

Software used for analysis is public and described in detail in the Methods section. Raw scripts and code are available upon reasonable request.

---

**Supplementary Table 1**

Patient clinical and demographic information for all 12 samples used for analysis.

**Supplementary Table 2**

Genes utilized for tumor core and leading edge annotation, identified through literature search.

**Supplementary Table 3**

Differentially expressed transcription factors between tumor core and leading edge spots. Table contains logFC and adjusted p-value information for all differentially expressed genes.

**Supplementary Table 4**

DEGs between tumor core and leading edge spots. Table contains logFC and adjusted p-value information for all differentially expressed genes.

**Supplementary Table 5**

DEGs factors between spatially deconvolved tumor core and leading edge cancer cells. Table contains logFC and adjusted p-value information for all differentially expressed genes.

**Supplementary Table 6**

Cell chat-predicted cell-cells signaling ligand receptor pairs and signaling strength (interaction probability). Table contains leading edge cancer cells, tumor core cancer cells, and leading edge ecm-myCAF cells as both sources and targets of signaling.

**Supplementary Table 7**

Differentially spliced genes identified with scVelo are ordered based on fit likelihood. Genes differentially spliced based on leading edge cancer cells, tumor core cancer cells, and transitory cancer cells are indicated.

**Supplementary Table 8**

Trimmed IC50 means for 220 drugs with IC50 values in at least 25 HPV negative HNSCC cell lines. Drugs are ordered based on decreasing trimmed IC50 mean value.

**Supplementary Video 1**

Dynamo vector field topology for tumor core cancer cells, leading edge cancer cells, and transitory cancer cells. Circular nodes are indicative of stable fixed points, half circles indicate saddle points. Black text colors represent absorbing points, red text colors represent emitting points, and blue text colors represent unstable points.

**Extended Data Fig. 1:**
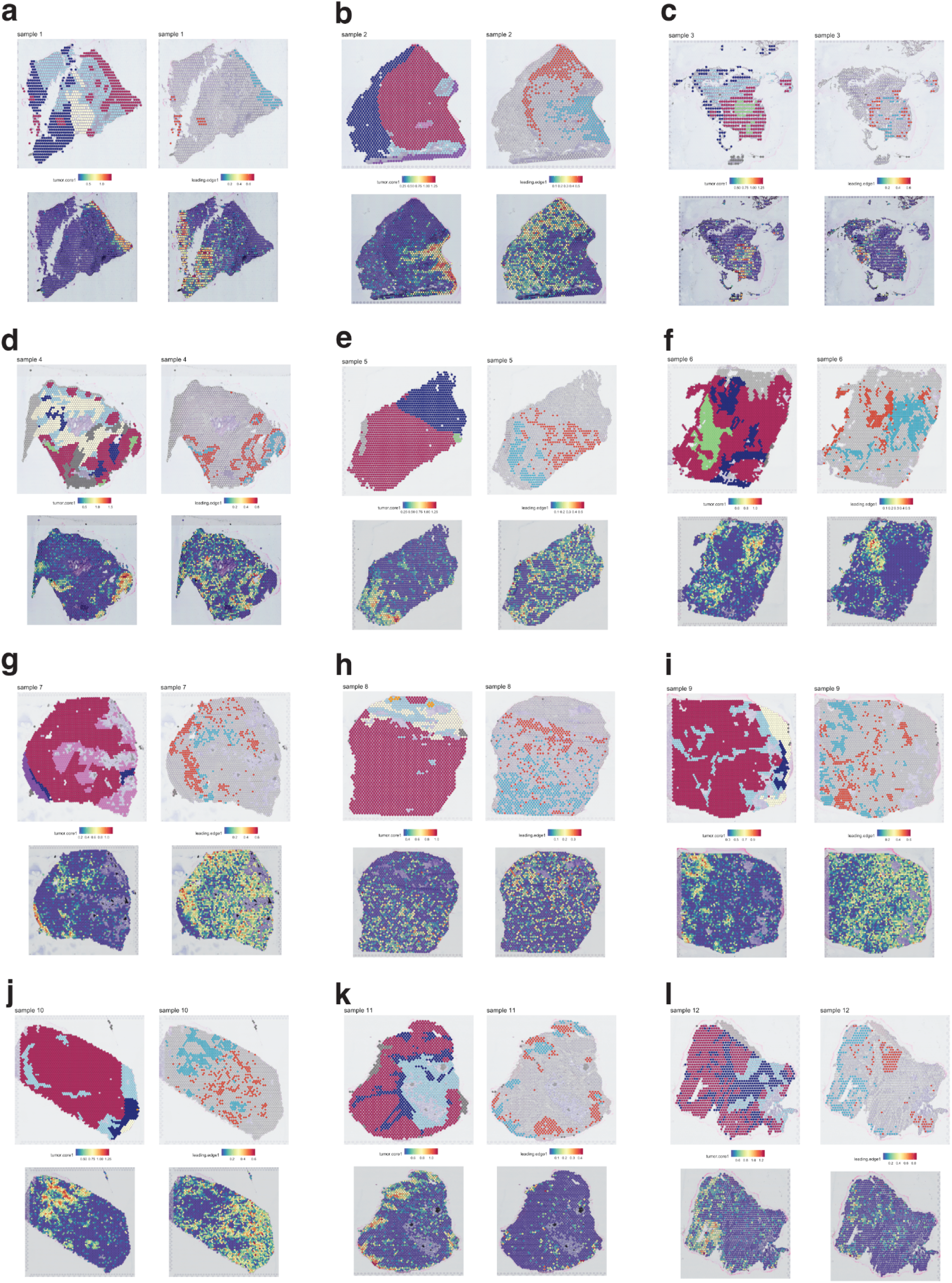
Annotation of tumor core and leading edge across all 12 tissue samples. **a-l**. Pathologist annotations, tumor core discovery geneset scores, leading edge discovery geneset scores, and final tumor core and leading edge annotations for samples 1-12. Geneset scores for the tumor core and leading edge are visualized for spots with a score greater than the first quartile and median respectively.

**Extended Data Fig. 2:**
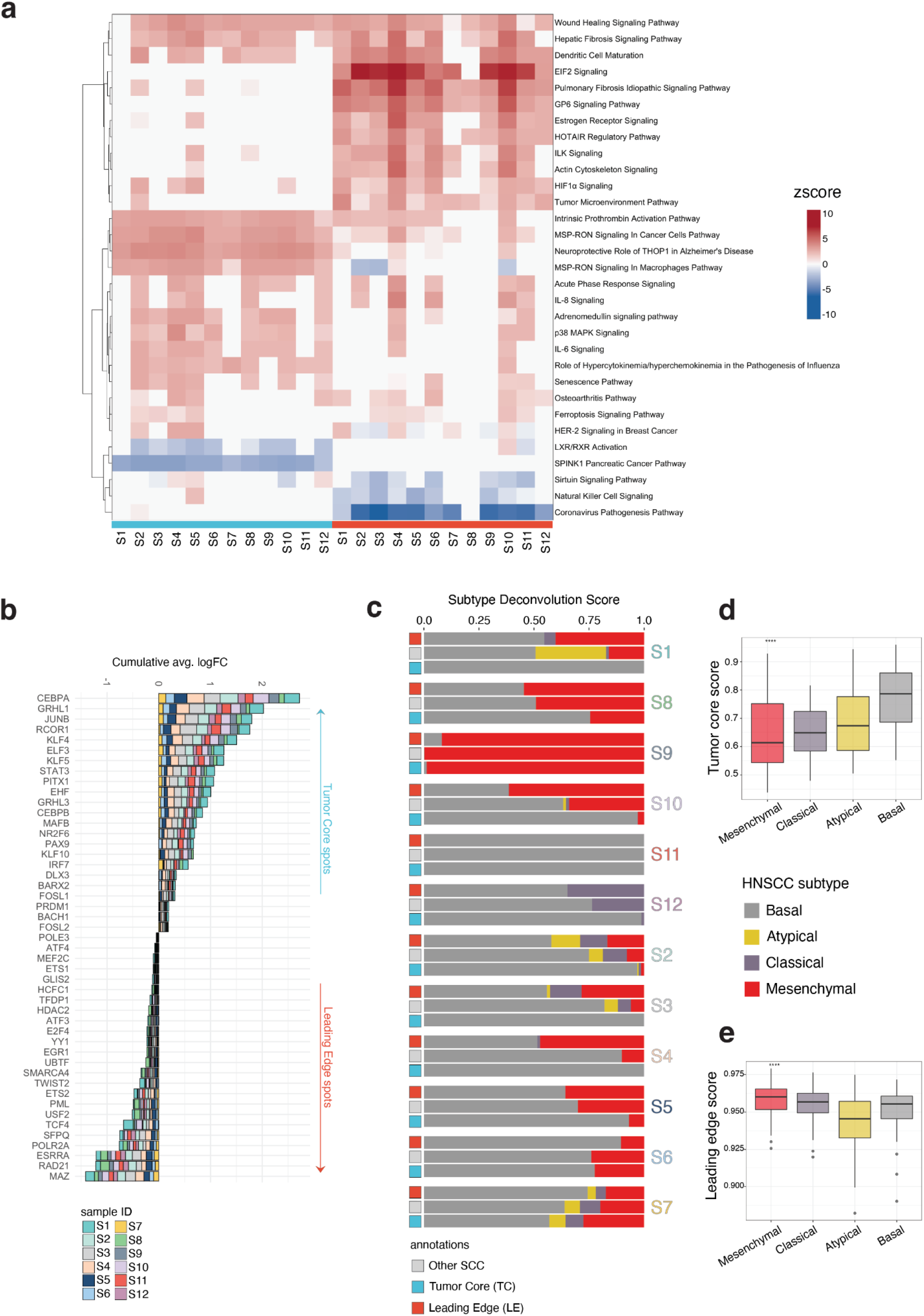
Core and edge tumoral regions are differentially regulated and not explained by HNSCC subtypes. **a**. Ingenuity Pathway Analysis pathway analysis plot visualizing predicted activation and deactivation of tumor core and leading edge pathways. Pathways are displayed if they are activated or deactivated across 10 samples and ordered based on similarity of z-score for each pathway across samples. **b**. Barplot displaying the cumulative average logFC for differentially expressed transcription factors between the tumor core and leading edge across more than 9 samples, inferred by SCENIC. **c**. Stacked bar plot visualizing HNSCC molecular subtype deconvolution scores for tumor core, leading edge, and other SCC spots. Deconvolution scores were obtained through integration and label transfer with 279 HNSCC bulk RNA sequencing samples labeled by subtype from the NCI TCGA. **d-e**. Bar plot displaying single sample geneset enrichment scores for the tumor core and leading edge genesets across TCGA HNSCC molecular subtype data in 279 HNSCC patients. Significance was compared between groups using a Kruskal-Wallis one-way analysis of variance test. *p < 0.05, **p < 0.01, ***p < 0.001, ****p < 0.0001.

**Extended Data Fig. 3:**
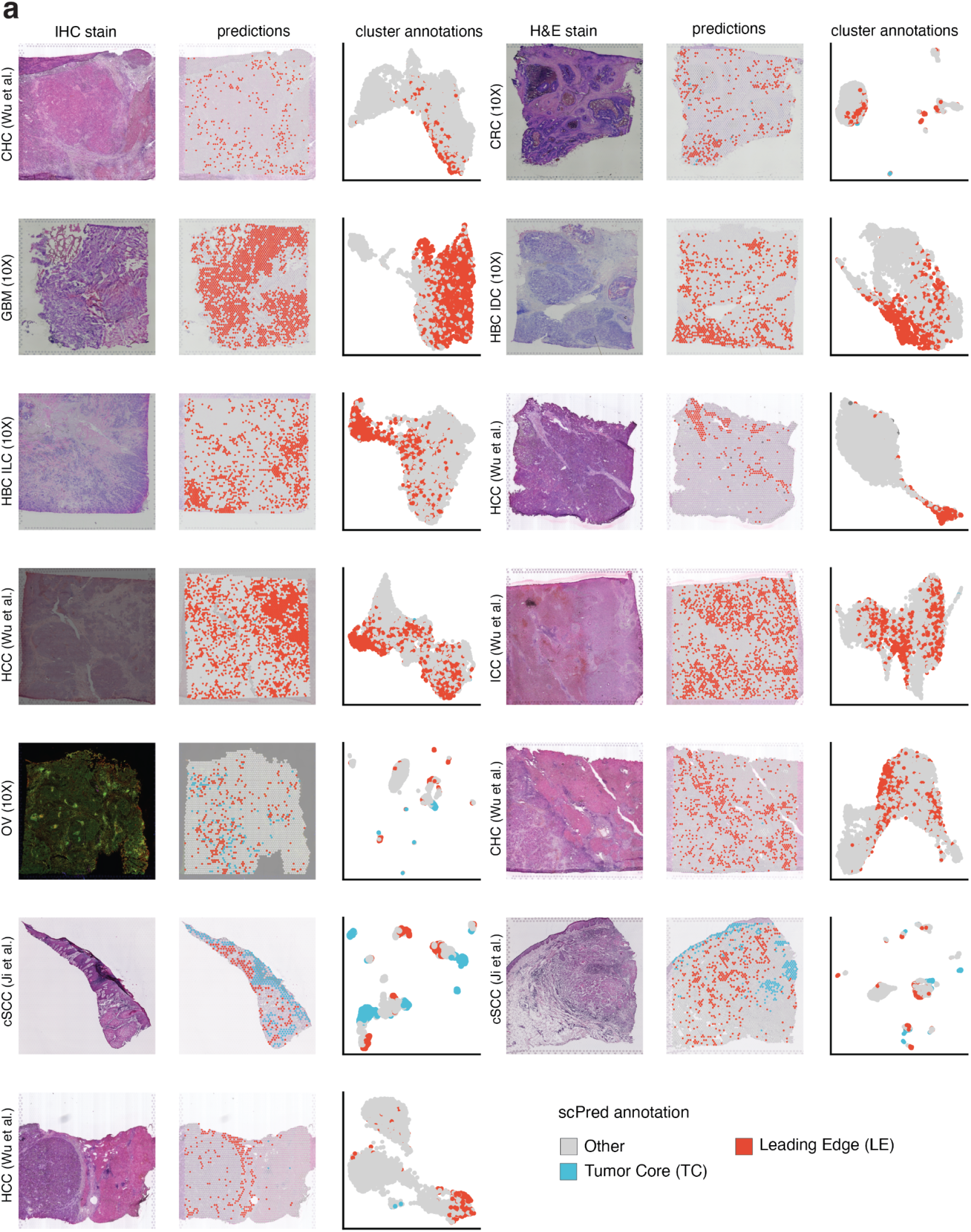
ScPred annotation of tumor core and leading edge states across 13 spatial datasets. **a**. Stained tissue section (left), scPred classification on stained tissue (middle), and a UMAP colored by scPred classification (right) for all 13 spatial transcriptomic datasets.

**Extended Data Fig. 4:**
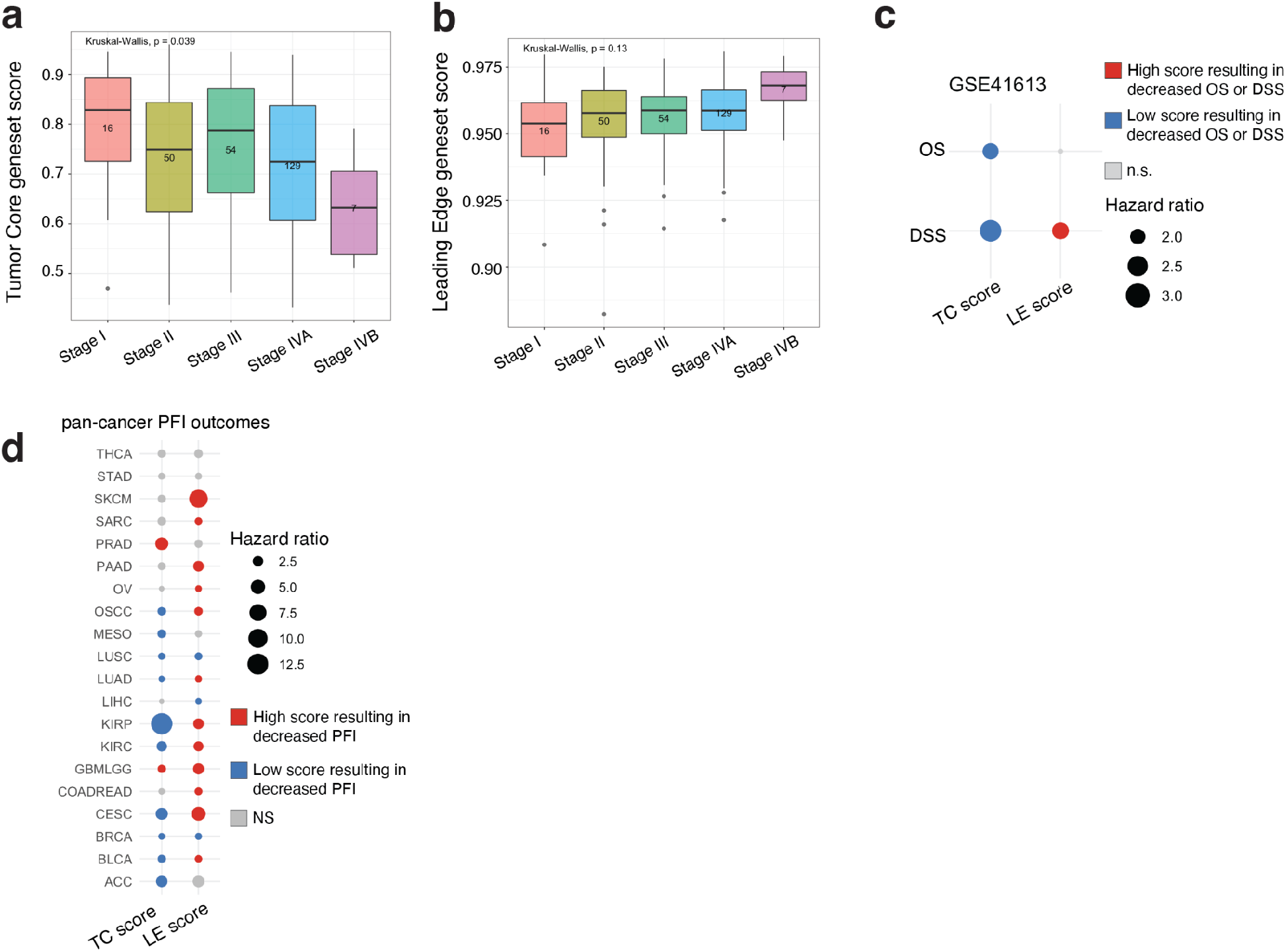
Core and edge regions are correlated to patient outcomes. **a-b**. Boxplots comparing tumor core and leading edge geneset single sample geneset enrichment score across pathological tumor stage. Significance was compared between groups using a Kruskal-Wallis one-way analysis of variance test. **c**. Overall survival and disease specific survival OSCC outcomes for tumor core and leading edge geneset enrichment scores in a validation dataset. P-values and hazard ratios displayed were calculated using a cox proportional-hazard regression. **d**. Progression free interval pan-cancer outcome for tumor core and leading edge geneset enrichment scores. P-values and hazard ratios displayed were calculated using a cox proportional-hazard regression.

**Extended Data Fig. 5:**
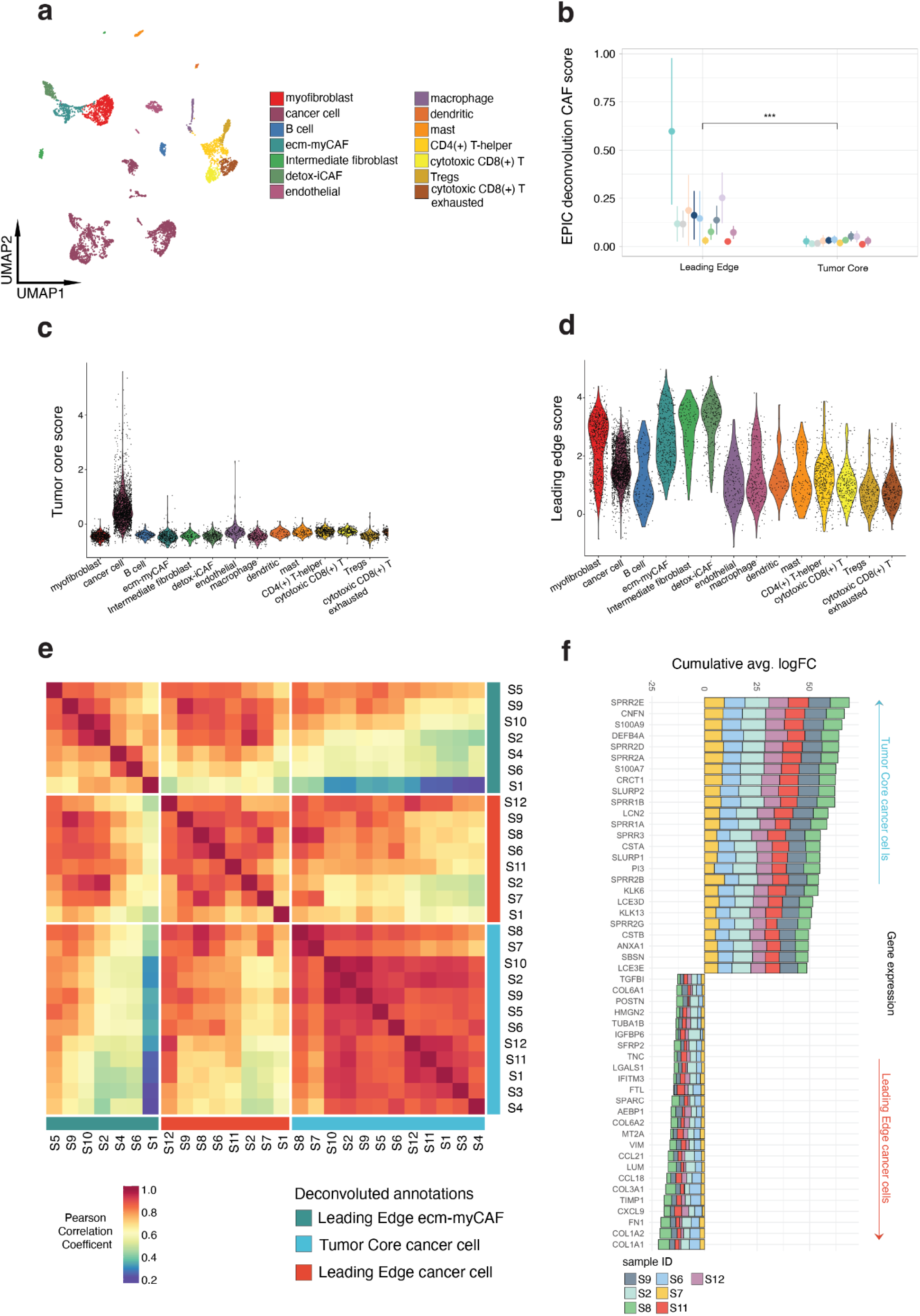
Characterization of spatially distinct cancer cells. **a**. Single cell UMAP plot of 23,691 cells from normal and malignant HNSCC samples colored by cell type identity. **b-c**. Violin plots of differentially expressed tumor core and leading edge geneset scores in the HNSCC single-cell RNA sequencing reference dataset across cell type identity. **d**. Comparison of EPIC CAF scores between leading edge and tumor core spots. Significance of differences between the tumor core and leading edge was determined through the use of a paired T test. Circles are representative of mean and lines are representative of IQR. All tested genesets had a significant p-value, *p < 0.05, **p < 0.01, ***p < 0.001, ****p < 0.0001. **e**. Whole transcriptome correlation heatmap of spatially deconvolved leading edge cancer cells, tumor core cancer cells, and leading edge ecm-myCAF annotations across all samples. Samples are ordered based on transcriptomic similarity. **f**. Stacked barplot displaying the cumulative average logFC for genes significantly (adj. p < 0.001) differentially expressed between spatially deconvolved tumor core and leading edge cancer cells.

**Extended Data Fig. 6:**
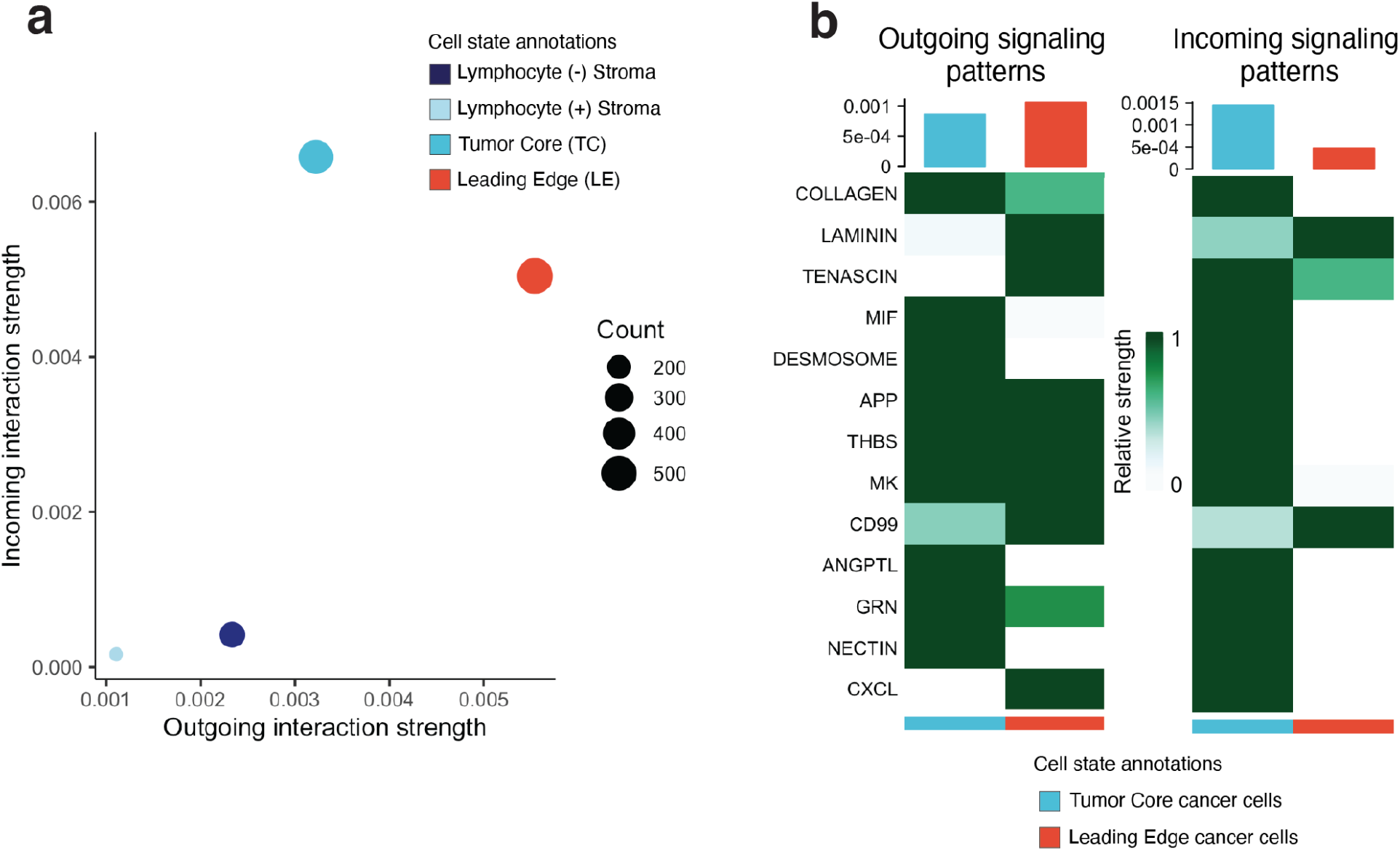
Cell signaling of stroma and cancer cell pathways. **a**. Scatterplot visualizing outgoing interaction strength and incoming interaction strength for tumor core spots, leading edge spots, and stromal spots. The number of inferred interactions is indicated by dot size. **b**. Outgoing and incoming signaling pathway strength for tumor core cancer cells and leading edge cancer cells. Signaling pathways are ordered based on strength of interaction.

**Extended Data Fig. 7:**
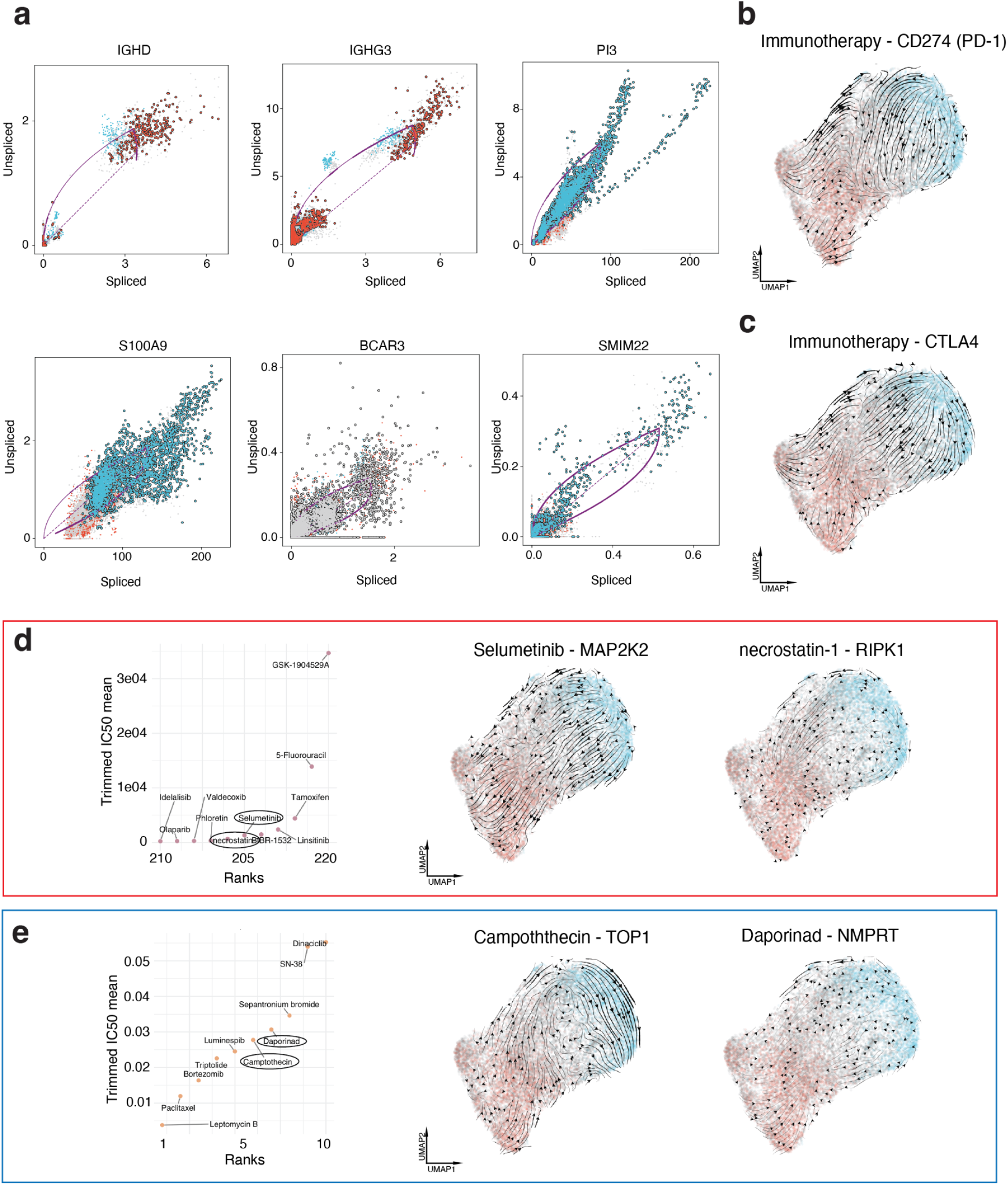
Splicing dynamics and drug responses of spatially unique cancer cells. **a**. Phase portraits showing the ratio of spliced and unspliced RNA ratios for top differentially spliced genes, purple lines depict splicing steady state. **b**. UMAP showing the resultant vector field following an in-silico perturbation of CD274 (PD-L1). **c**. UMAP showing the resultant vector field following an in-silico perturbation of CTLA4. **d**. Rank ordered dotplot for the 10 highest trimmed IC50 means. The fourth and fifth highest IC50 anticancer drugs and their genetic molecular targets are highlighted (left). UMAP showing the resultant vector field following in silico perturbations of drug targets (center, right). **e**. Rank ordered dotplot for the 10 lowest trimmed IC50 means. The fourth and fifth lowest IC50 anticancer drugs and their genetic molecular targets are highlighted (left). UMAP showing the resultant vector field following in silico perturbations of drug targets (center, right).

